# Host diversity and behavior determine patterns of interspecies transmission and geographic diffusion of avian Influenza A subtypes among North American wild reservoir species

**DOI:** 10.1101/2021.09.29.462321

**Authors:** Joseph T. Hicks, Kimberly Friedman, Xueting Qiu, Do-Kyun Kim, James E. Hixson, Scott Krauss, Richard J. Webby, Robert G. Webster, Justin Bahl

## Abstract

Wild birds can carry avian influenza viruses (AIV), including those with pandemic or panzootic potential, long distances. Even though AIV has a broad host range, few studies account for host diversity when estimating AIV spread. We analyzed AIV genomic sequences from North American wild birds, including 303 newly sequenced isolates, to estimate interspecies transmission and geographic diffusion patterns among multiple co-circulating subtypes. Our results show high transition rates within Anseriformes and Charadriiformes, but limited transitions between these orders. Patterns of interspecies transmission were positively associated with breeding habitat range overlap, and negatively associated with host genetic distance. Distance between regions (negative correlation) and summer temperature at origin (positive correlation) were strong predictors of diffusion. Taken together, this study demonstrates that host diversity and ecology can determine evolutionary processes that underlie AIV natural history and spread. Understanding these processes can provide important insights for effective control of AIV.

## INTRODUCTION

Avian influenza viruses (AIV) are globally distributed pathogens maintained within wild waterfowl (order Anseriformes) and shorebirds (order Charadriiformes) (1). Despite being largely asymptomatic within wild birds, AIV provide cause for global concern as sources of influenza A viral diversity for domestic avian and mammalian hosts (2). AIV hemagglutinin (HA) subtypes H5 and H7 have repeatedly evolved into highly pathogenic viruses in domestic poultry causing devastating loss (3). Furthermore, all modern pandemic influenza viruses contain gene segments of avian origin, suggesting reassortment with avian viruses plays a crucial role in pandemic emergence (4). The segmented genome is an important characteristic of influenza viruses because it facilitates continual reassortment and promotes diversity of AIV within wild avian populations (1,5,6). AIV segments are habitually interchanging, existing as functionally equivalent arrangements (5). Due to the unlinked nature of the AIV genes, each segment can be considered as an independent hereditary particle with its own evolutionary history (7).

Understanding the host behavior and environmental drivers of AIV susceptibility and dispersal remain a top priority for avian influenza surveillance, but the vast array of susceptible host species and ecological variables hampers the prediction of AIV emergence and incidence (8). Surveillance data and spatial analysis have begun to assess the association between avian influenza prevalence and environmental variables, including land use (9,10), temperature measures (9,11,12), altitude (10), distance to water (10), and precipitation (11). Fewer studies have assessed the impact of host characteristics on the prevalence of AIV within individual avian species although migration distance, habitat water salinity, and surface foraging methods have been implicated as important predictors in one such study (13). Sequence data acquired by viral surveillance provide further information to understand AIV dynamics. Because viral evolution, host ecology, and environmental factors necessarily interact, phylogenetic studies can help elucidate the paths of AIV dispersal (Figure 1). For example, previous phylogeographic analysis (7) of AIV within North America provided evidence that migratory flyways are not as strong a barrier to viral dispersal as previously believed (14).

**Figure 1.**
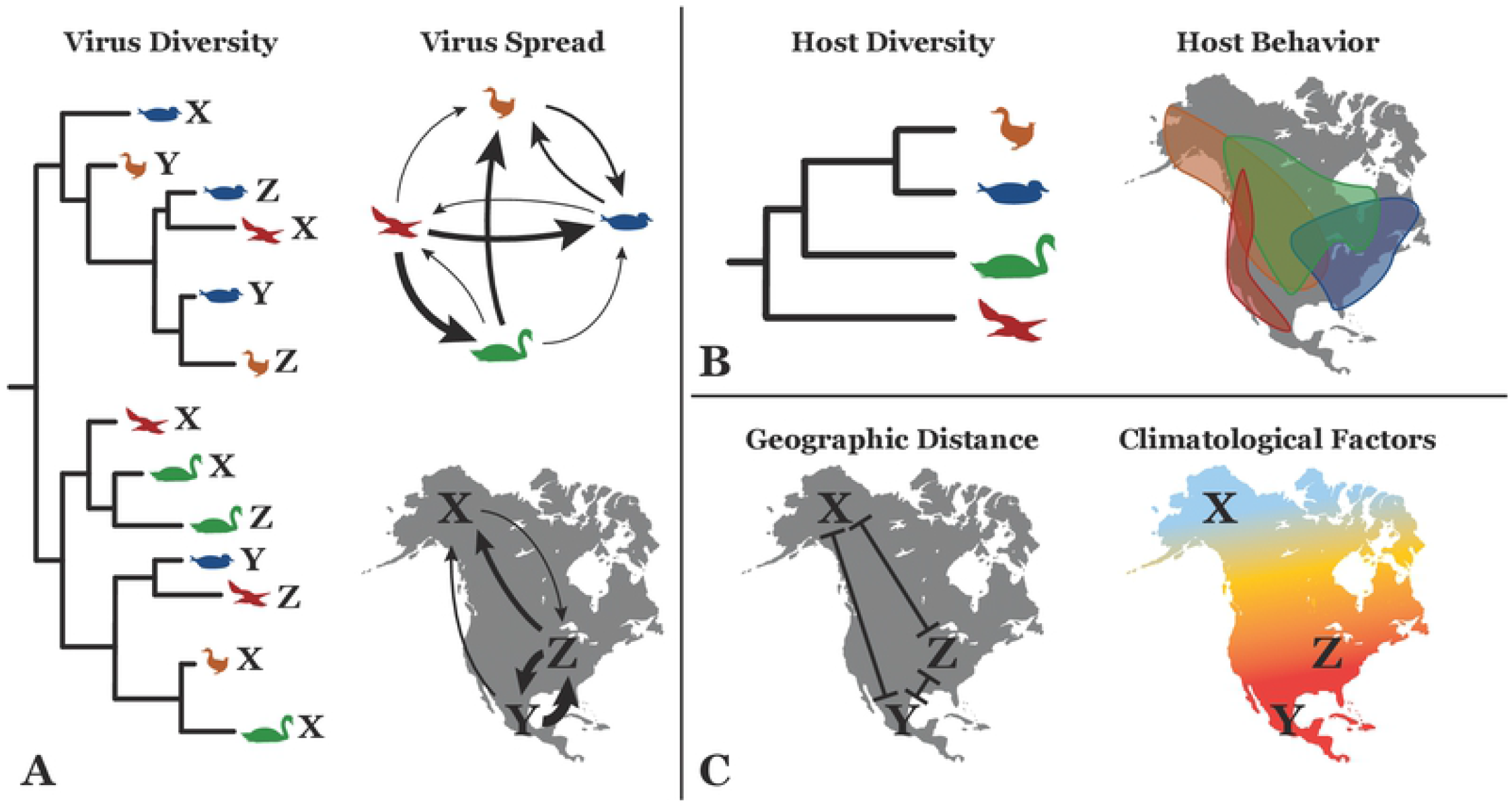
Interaction of viral evolution, host ecology and the environment. Viral genetic sequences contain information regarding virus evolution and diversity (A). Because their evolution occurs at a rapid pace, evolutionary patterns can be used in conjunction with location and species data to infer rates of viral dispersal among sampled geographic regions and host species. Many factors may influence observed virus transmission and spread. For instance, host factors (B) such as relatedness of host species and overlap of habitat distributions may be associated with viral transitions between host species. Further, environmental factors (C) may also play a role in the spatial diffusion of the virus. By incorporating viral, host and environmental information into computational models, the impact of host and environmental characteristics on virus spread can be estimated.

Although phylogenetic studies to date have been able to interrogate the impact of broad ecological patterns such as migratory flyways and interhemispheric viral exchange, few incorporate characteristics of the location or host from which the virus was sampled. Prevalence studies include these characteristics into regression and spatial models, but are limited due to the long-distance migration of wildlife hosts. Generalized linear models (GLM) implemented within a Bayesian phylogenetic framework have made it possible to include environmental and ecological covariates into phylogenetic models (15,16). This allows the simultaneous inference of viral transition rates among specified traits (i.e., hosts or locations) and their association with covariates that may drive viral movement. This approach has been adapted to investigate the role of anthropogenic and environmental variables on the diffusion of avian influenza within China (17) as well as of avian influenza subtype H9N2 on a global scale (18). The GLM has also been used to uncover the impact of host behavior on the dispersal of rabies virus among bat species (19). Previous analyses have also demonstrated the importance of environmental transmission on AIV prevalence and evolution (20,21), suggesting ecological factors may influence AIV transmission. Understanding how avian host characteristics and environmental variables impact zoonotic transmission and geographic dispersal will be key to identify surveillance priorities among species and locations. In the presented analysis, using an extensive publicly-available dataset of multiple AIV subtypes collected from North American wild birds supplemented with newly sequenced surveillance samples, we implemented the GLM to assess the impact of ecological and environmental characteristics on the dispersal of AIV across the North American continent and among frequently sampled Anseriformes and Charadriiformes.

## RESULTS

### Summary of Newly Sequenced Data

Supplementary Table S1 describes the characteristics of 303 newly sequenced AIV isolates, which originated from samples collected from wild birds between 2003 and 2016 in Delaware Bay, New Jersey, United States (86.5%) and Alberta, Canada (13.5%). All sequenced samples from Alberta were exclusively of waterfowl origin (order Anseriformes). Delaware Bay samples originated almost exclusively from shorebirds (order Charadriiformes), except for a single Canada goose (*Branta canadensis*) sample. Among all newly sequenced viral isolates, most (60.4%) were isolated from samples collected from the ruddy turnstone (*Arenaria interpres*), a migratory shorebird of the wader family with near global distribution and intercontinental migration patterns. Nine samples were found to be co-infected with avian paramyxovirus and were excluded from further analysis. The most frequently isolated hemagglutinin (HA) subtype was H10 (27.4%), followed by H12 (18.8%) and H3 (9.2%). Most HA subtypes were collected in Delaware Bay, including H1, H3, H5, H6, H7, H8, H9, H10, H11, H12, H13, and H16. Only H4 was exclusively isolated from Alberta. The most frequent neuraminidase (NA) subtypes were N5 (20.1%), N7 (13.9%), and N8 (13.5%). All but two NA subtypes were isolated from both Delaware Bay and Alberta; N3 and N9 were only recovered from Delaware Bay.

### Evolutionary Comparison Between Segments and Subtypes

The newly sequenced data were aligned with publicly available sequences and subsampled in two methods to help address sampling biases in surveillance: a phylogenetic diversity-based analysis method (PDA sample) and a simple stratified random sample method (stratified sample). Evolutionary models were constructed separately for each gene segment; HA, NA, and NS segment datasets were further subdivided by subtype or allele. Because the PDA sample maintains the total genetic diversity of the original sample, the evolutionary parameters of the PDA sample are discussed here. In general, the two samples produced similar comparative relationships of evolutionary parameters among the analyzed gene segments; however, the stratified sample had consistently lower molecular clock rates and effective population sizes compared to the PDA sample (Supplementary Figure S1). Compared with the HA and NA surface proteins, the internal gene segments tended to have older times to the most recent common ancestor (TMRCA) (Supplementary Figure S1A; Supplementary Table S2), except the included sequences of the NS gene B allele, which shared a common ancestor around 1965 (95% Highest Posterior Density Bayesian Credibility Interval (HPD) 1958.5 – 1969.7). HA and NA genes tended to have a TMRCA within the mid-to latter-half of the twentieth century, although this pattern deviated for H3 and N3, which had TMRCA older than other HA and NA subtypes (H3: 1929.0, 95% HPD 1900.0 – 1948.5; N3: 1895.6, 95% HPD 1830.8 – 1944.7). The uncertainty of the N3 subtype TMRCA is most likely due to the lack of older sequences which would help calibrate the divergence time of apparent Eurasian-origin viruses that circulate in North America.

As compared to the HA and NA surface proteins which contend with greater selection pressure, the internal gene segments tended to have slower evolutionary rates as measured by the mean substitution rate of the uncorrelated relaxed molecular clock (Supplementary Figure S1B; Supplementary Table S2). The two NS alleles had differing substitution rates with the A allele evolving faster (3.4×10^-3^ substitutions/site/year; 95% HPD 3.0×10^-3^ – 3.7×10^-3^) compared to the B allele (2.6×10^-3^ substitutions/site/year; 95% HPD 2.3×10^-3^ – 2.9×10^-3^). In comparison to the internal gene segments, the HA and NA surface proteins were estimated to have more variable substitution rates with median rates ranging from 2.5×10^-3^ (H3) to 5.8×10^-3^ (H7) substitutions/site/year. Six of the eight analyzed HA subtypes had estimated median substitution rates above 3.8×10^-3^ substitutions/site/year. In contrast, H3 and H4 were estimated to have much slower substitution rates at 2.5×10^-3^ (95% HPD 2.2×10^-3^ – 2.7×10^-3^) and 2.8×10^-3^ (95% HPD 2.6×10^-3^ – 3.1×10^-3^) substitutions/site/year, respectively. The NA subtype with the fastest substitution rate was N7 (5.0×10^-3^ substitutions/site/year; 95% HPD 4.4×10^-3^ – 5.6×10^-3^).

Internal gene segments were estimated to be sustained by a much larger effective population size as compared to the surface proteins (Supplementary Figure S1C; Supplementary Table S2). NS alleles differed from the remaining internal gene segments, with median effective population sizes around half that of PB2, PB1, PA, NP, and MP segments. Similarly, the effective population sizes of the various HA and NA subtypes were considerably lower than that of the non-subdivided internal gene segments. Because the genetic diversity of the NS, HA, and NA gene segments was divided between datasets, lower population sizes are needed to explain the observed viral circulation. Variation among HA and NA subtypes was also noted. Most HA subtypes were estimated to have median effective population sizes below 20, but those of H3 and H4 were substantially larger (H3: 99.3, 95% HPD 87.2 – 112.5; H4: 78.0, 95% HPD 68.3 – 88.2). The effective population sizes of the NA subtypes also varied considerably with median sizes ranging from 8.5 (N7) to 85 (N8).

### Discrete Trait Diffusion Models

Two discrete trait diffusion models were estimated for each of 22 gene segment or subtype datasets to assess how AIV disperses among host species and geographic regions of North America. North American regions were categorized into eight Canadian provinces and territories (Alberta, British Columbia, New Brunswick, Newfoundland and Labrador, Nova Scotia, Ontario, Prince Edward Island, and Quebec), ten United States climate regions (Alaska, Midwest, Northeast, Northwest, Ohio Valley, Northern Rockies and Plains, South, Southeast, Southwest, and West), one Mexican state (Sonora), and Guatemala. Represented host species were of the taxonomic orders Anseriformes (waterfowl) and Charadriiformes (shorebirds). The 16 Anseriformes species included American black duck (*Anas rubripes*), bufflehead (*Bucephala albeola*), blue-winged teal (*Anas discors*), Canada goose (*Branta canadensis*), cinnamon teal (*Anas cyanoptera*), emperor goose (*Anser canagicus*), gadwall (*Mareca strepera*), greater white-fronted goose (*Anser albifrons*), green-winged teal (*Anas crecca*), mallard (*Anas platyrhynchos*), northern pintail (*Anas acuta*), redhead (*Aythya americana*), ring-necked duck (*Aythya collaris*), northern shoveler (*Anas clypeata*), snow goose (*Anser caerulescens*), and American wigeon (*Anas americana*). Five species of Charadriiformes were represented among the host models: glaucous-winged gull (*Larus glaucescens*), laughing gull (*Leucophaeus atricilla*), red knot (*Calidris canutus*), ruddy turnstone (*Arenaria interpres*), and sanderling (*Calidris alba*). Henceforth, all host species will be referenced using common names.

The distribution of hosts and geographic regions were similar among the internal gene segments by both PDA and stratified subsampling strategies (Supplementary Table S3; Supplementary Table S4). Subsampling strategy had an effect on the temporal distribution of host and geographic region variables; proportions were more consistent between 2005 and 2016 in the stratified sample compared with the PDA sample (Supplementary Figures S2 & S3). Within both the PDA and stratified samples, Alaska, the Ohio Valley, and the Northeast were the most frequently represented regions between 2005 and 2016 among the internal genes. Regional distribution varied considerably across HA and NA subtypes. Within the PDA sample, the most frequently represented regions’ HA subtypes included Alaska (H3: 30.5%, H4: 20.5%), Northeast (H1: 20.7%, H10: 31.1%, H11: 20.0%), South (H7: 23.7%), and West (H5: 20.5%, H6: 28.1%, H11: 20.0%). The most frequently represented regions among NA subtypes in the PDA sample included Alaska (N3: 18.3%, N6: 25.9%, N8: 24.5%), Midwest (N2: 20.3%, N9: 20.3%), and Northeast (N1: 19.2%, N7: 25.2%). Although host species distribution differed among gene segments and subtypes, all shared mallard as the most frequently sampled avian species (28.1 – 52.7%). The stratified sample attempted to counteract the oversampling of mallards resulting in lower percentages of these hosts within the sample (20.6 – 35.7%). The stratified method also tended to increase the frequency of the more sparsely represented regions and hosts within the models.

Asymmetrical diffusion models allow directionality to be inferred so that each viral transition is characterized by a source (i.e., origin of the virus) and a sink (i.e., destination). Because the rates of transition between two locations, for instance, are asymmetrical, the transition rate from location A to location B is permitted to differ from the rate in the opposing direction (from location B to A). Across all gene segments, the highest rate of transition between host species was 19.6 transitions/year (95% HPD 16.0 – 23.6) of the PB1 gene segment from mallards to blue-winged teals within the PDA sample (Supplemental Figure S4). In the PDA sample, the N8 transition rate from mallards to blue-winged teals was highest among the NA subtypes at 13.3 transitions/year (95% HPD 9.3 – 17.7). In contrast, among HA subtypes, the reverse transition (that is, blue-winged teals to mallards) within the PDA H4 model had the highest rate (16.1 transitions/year; 95% HPD 10.9 – 21.4). Mallards were supported as the source of AIV across all gene segments and subtypes for green-winged teals and northern shovelers within the stratified sample. These rates were also supported in the PDA sample in all gene segments and subtypes except N6. The rates from mallards to blue-winged teals were also supported among all gene segments and subtypes except H7 in both samples. Within the PDA sample, American black ducks, blue-winged teals, Canada geese, greater white-fronted geese, ring-necked ducks and snow geese were only supported to receive viral diversity from mallards. In contrast, only American black ducks and Canada geese exclusively received virus from mallards in the stratified sample. Similarly, ruddy turnstones were the exclusive source of viral diversity for laughing gulls and sanderlings in both samples, as well as red knots in the stratified sample.

A single host diffusion model was also jointly estimated across all internal gene segments, all HA subtypes, and all NA subtypes (Figure 2, Supplementary Figure S5). For each joint host model, the highest transition rate occurred from blue-winged teals to mallards (PDA internal gene model: 40.7 transitions/year, 95% HPD 32.5 – 49.2; PDA HA model: 22.0 transitions/year, 95% HPD 16.6 – 27.5; Stratified NA model: 22.6 transitions/year, 95% HPD 14.2 – 31.7; Tables S5, S6 & S7). In all three joint host models, all species were supported as receiving virus from at least one other species, except snow geese within the NA models. Not all species acted as a source, however. Cinnamon teals, gadwalls, and red knots were included in all three joint host models, but none were supported to contribute AIV genetic diversity to any other host species. In addition, Canada geese, emperor geese, ring-necked ducks, snow geese, laughing gulls, and sanderlings, which were only included in the internal gene and NA models, were also not supported as viral sources. A marked difference between the PDA and stratified samples can be noted in this regard. Whereas green-winged teals were not supported as a source of virus for any other species within the PDA internal gene segment model, the stratified sample estimates green-winged teals as the source of viral genetic diversity for nine other avian species within the internal gene model. This provides evidence that sampling methods can influence discrete trait diffusion model results.

**Figure 2.**
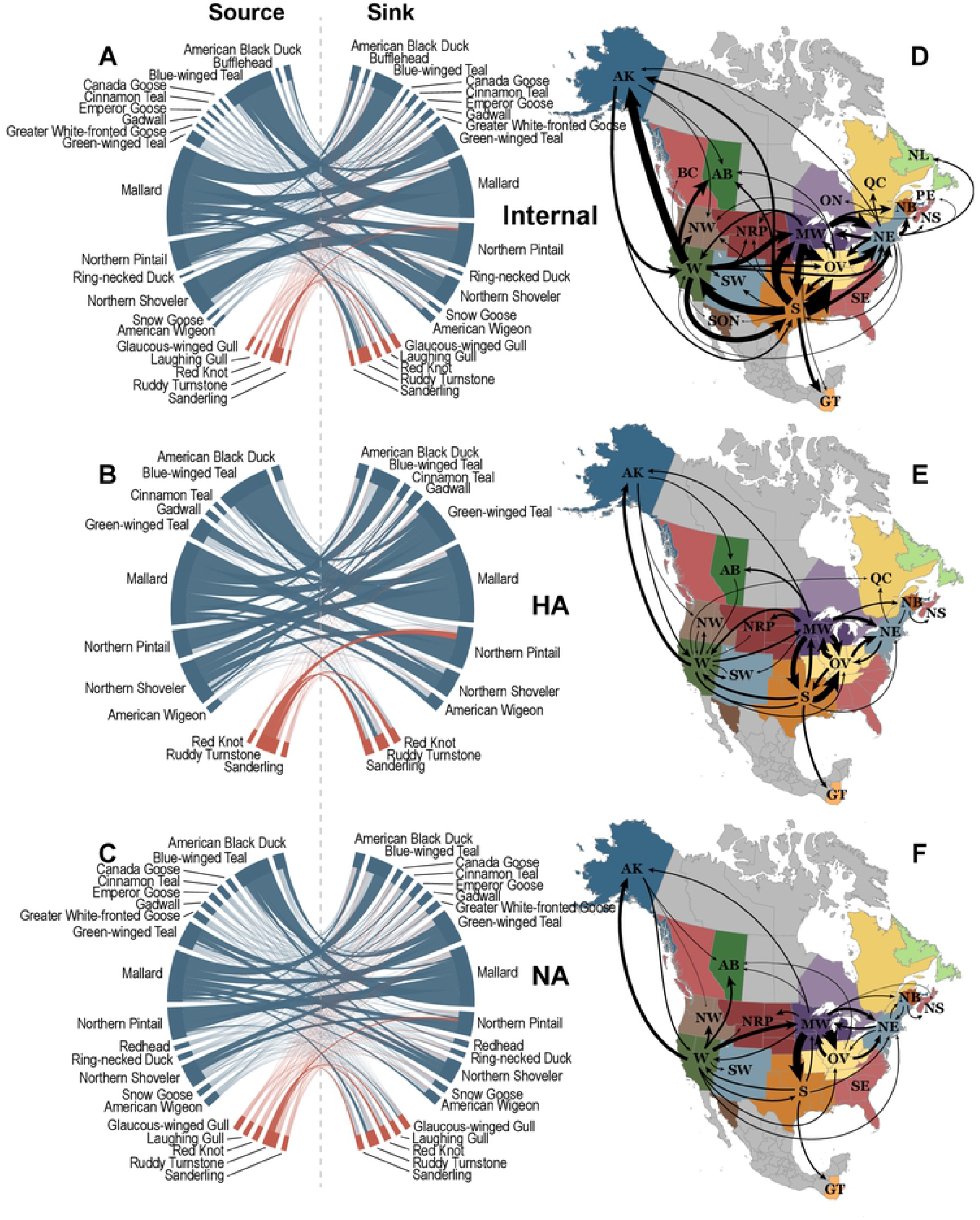
Discrete trait diffusion models of North American avian influenza using a sample of genetic sequences based on phylogenetic diversity. Host models (left) are presented for combined internal gene segment (A), hemagglutinin gene subtype (B), and neuraminidase gene subtype (C) models. Source host species on the left of the chord diagrams contribute viral diversity to sink host species on the right. The magnitude of the viral transition rate is proportional to the width of the band, and statistically supported rates are darkened. Bands are colored by the host order of the source species (Charadriiformes – red; Anseriformes – blue). Similarly, geographic models (right) are summarized for combined internal gene segment (D), hemagglutinin gene subtype (E), and neuraminidase gene subtype (F) models. Arrow width is proportional to the magnitude of the transition rate, and only statistically supported rates are displayed. (AK – Alaska, AB – Alberta, BC – British Columbia, GT – Guatemala, MW – Midwest, NB – New Brunswick, NE – Northeast, NL – Newfoundland and Labrador, NRP – Northern Rockies and Plains, NS – Nova Scotia, NW – Northwest, OV – Ohio Valley, PE – Prince Edward Island, QC – Quebec, S – South, SE – Southeast, SON – Sonora, SW – Southwest, W – West)

Among the North American regional models, the highest transition rate was observed from the Ohio Valley to the South within the PA gene segment of the PDA sample at a rate of 23.3 transitions/year (95% HPD 17.6 – 29.0; BF = 73,262) (Figure S6). In the PDA sample, the N6 model showed the highest transition rate of the NA subtypes with a Midwest to South transition rate of 11.4 transitions/year (95% HPD 7.5 – 15.7; BF = 46,210). Within the stratified sample, the highest transition rate among HA subtypes occurred from the Midwest into the Ohio Valley in the H4 model at 14.9 transitions/year (95% HPD 9.9 – 20.1, BF = 46,210). No single transition rate was supported across all gene segments or subtypes. The internal gene segments and the HA and NA subtype models differed in regard to support for the Northeast region of the United States as a source of AIV for other North American regions. Across the HA and NA subtypes, there is only sporadic support for the Northeast as a source of AIV, with only three rates among the subtypes supported in the PDA sample, and four supported in the stratified sample. In contrast, each internal gene segment model within the PDA sample has at least six rates in support of the Northeast as a source. Support for a Northeastern source is less consistent across the stratified sample internal gene segments: while nine rates are supported in the PB2 model, no rates in the PB1 model are supported. The internal genes further differ between PDA and stratified samples in terms of their support for New Brunswick as a viral source. No New Brunswick source rates are supported within the PDA internal gene segment models, yet 16 rates are supported in the stratified models among four sink regions (Northeast, Nova Scotia, Ohio Valley, and Prince Edward Island).

Among the three joint models, the internal gene model has the largest number of decisively supported transition rates between regions (Figure 2, Supplementary Figure S5). The highest rate among the internal genes occurred from the South to the Ohio Valley (PDA sample: 48.5 transitions/year, 95% HPD 42.5 – 55.0; Table S8). The highest transition rate among both HA and NA models occurred from the Midwest to the Ohio Valley (stratified HA: 17.8 transitions/year, 95% HPD 12.7 – 23.4; PDA NA: 21.3 transitions/year, 95% HPD 16.7 – 26.2; Tables S9 & S10). Similar patterns can be observed across the three models. For instance, due to their frequent support and large transition rates, the West, Midwest, South, and Ohio Valley all appear to be important regions in the dispersal of AIV across the North American continent. Furthermore, while most decisively supported rates are between neighboring regions, longer distance transitions are also observed in all three models, including between the West and Alaska, the South and Guatemala, and the West and the Ohio Valley. Many supported rates also align with an East-West axis, suggesting viral exchange across migratory flyways.

### Generalized Linear Model

The discrete trait diffusion models were extended with a GLM to evaluate the impact of host and geographic ecological characteristics on AIV dispersal among host species and geographic regions within North America. Table 1 summarizes host species and regional characteristics included in the GLM. Genetic distance of host species ranged widely, from 0.3 to 196.7 million years (Figure 3). As expected, the host phylogeny is composed of two main clades, Anseriformes and Charadriiformes, which diverged between 86.6 and 110.2 million years ago. Included Charadriiformes species shared a most recent common ancestor between 46.0 and 64.2 million years ago, whereas Anseriformes species diverged into familial clades of Anatidae (ducks) and Anser (geese) between 10.0 and 31.4 million years ago. All nodes within the phylogeny were highly supported (posterior probability > 0.98) except for the ancestral node of the green-winged teal and Northern pintail (posterior probability = 0.67). Host habitat distributions tended to have greater percentage of overlap during the nonbreeding winter months versus the breeding summer months. Migration propensity and distance were calculated based on distribution maps and varied widely among included species. On average, 84% of a species’ breeding season distribution was considered migratory as opposed to resident. The average distance between North American geographic regions was 2,716 km, and geographic regions overlapped with 47% and 48% of the breeding and nonbreeding distributions, respectively. Regions tended to have greater precipitation, lower humidity, and more lush vegetation during the summer months compared to the winter months. Variables such as host genetic relatedness, habitat overlap, and geographic distance reflect the relationship between two variables, whereas the remaining ecological variables summarize aggregate measurements. For this reason, the relational variables were only included in the GLM once, but the remaining characteristics were each included twice to capture directionality of the viral transition rate. For instance, the average temperature during the summer months was included twice to assess if the summer temperature of the source region was associated with viral transition or if the summer temperature of the sink region impacted viral transition.

**Figure 3.**
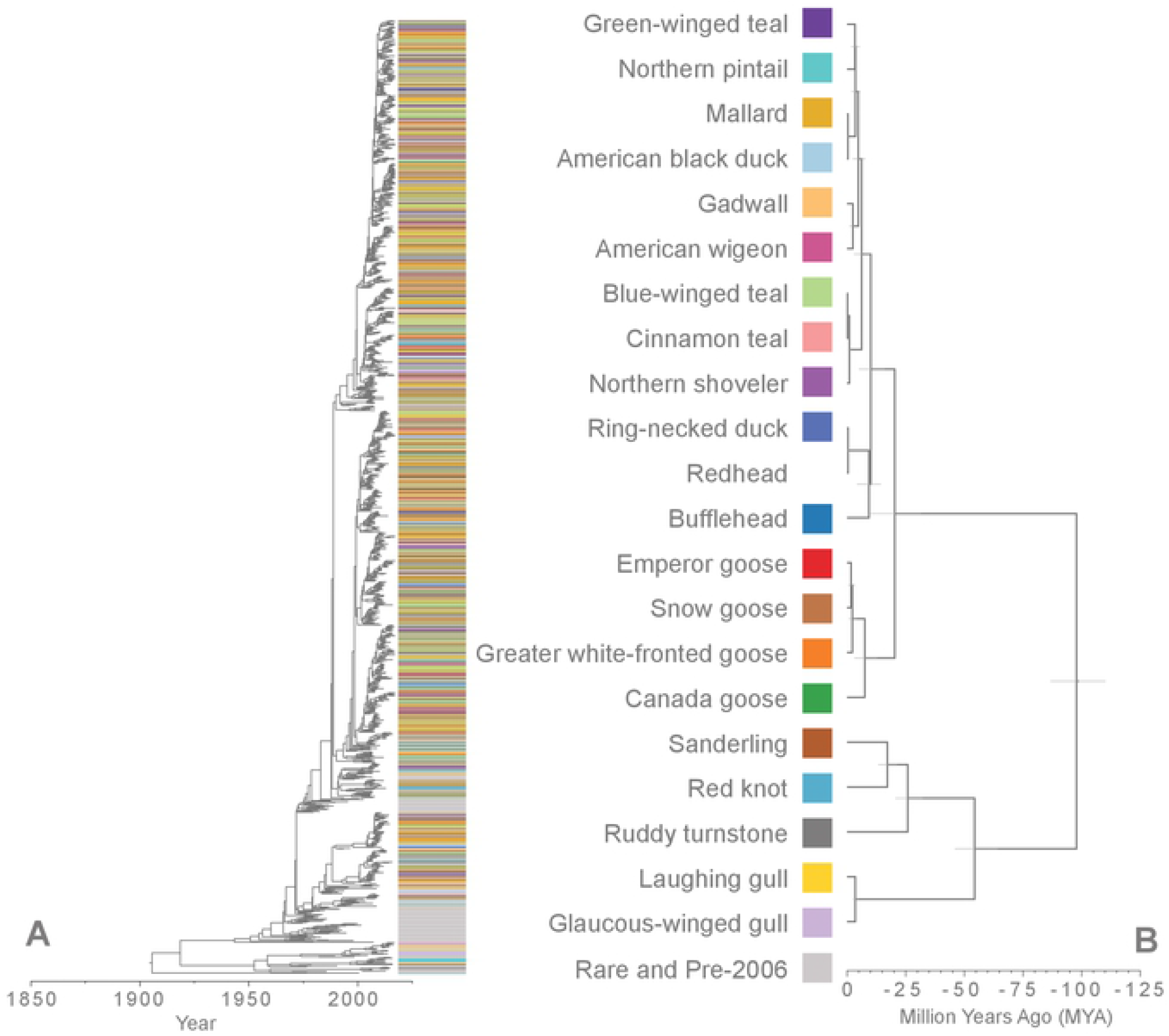
Viral and host phylogenetic diversity of North American AIV. (A) Estimation of the phylogenetic history of the PB2 AIV gene segment within North American wild birds. Color bands at the tips of the tree denote the host species distribution. This is contrasted with the phylogenetic history of the avian host species included in this analysis (B). Light gray node bars represent the 95% highest posterior density of the node height. The redhead species was not categorized in the internal gene segment models and is therefore not included.

**Table 1.** Summary of host and geographic variables used to inform the Bayesian discrete diffusion generalized linear model describing avian influenza virus dispersal among North American wild birds.

|  | Mean | Standard Deviation | Range |
| --- | --- | --- | --- |
| Genetic Distance (Myr) | 92.9 | 83.4 | 0.3 – 196.7 |
| Summer Distribution Overlap (%) | 26.6 | 32.0 | 0.0 – 99.99 |
| Winter Distribution Overlap (%) | 40.9 | 32.2 | 0.0 – 99.6 |
| Breeding Latitude | 56.4 | 11.7 | 28.0 – 76.7 |
| Nonbreeding Latitude | 34.1 | 8.3 | 21.6 – 56.6 |
| Migration Distance (km) | 2469.2 | 1447.9 | 473.8 – 5651.8 |
| Migration Propensity (%) | 84.4 | 22.1 | 18.1 – 100.0 |
| Geographic Distance (km) | 2716.9 | 1342.2 | 138.5 – 7087.5 |
| Summer Proportion (%) | 46.7 | 20.3 | 0.0 – 85.7 |
| Winter Proportion (%) | 48.1 | 28.2 | 0.0 – 90.5 |
| Summer Temperature (C) | 19.7 | 5.1 | 11.0 – 28.9 |
| Winter Temperature (C) | -2.7 | 9.4 | -15.6 – 19.7 |
| Summer Precipitation (kg/m<sup>2</sup>) | 2.7 | 1.4 | 0.3 – 7.5 |
| Winter Precipitation (kg/m<sup>2</sup>) | 1.9 | 0.9 | 0.4 – 3.4 |
| Summer Humidity (%) | 66.9 | 15.7 | 31.4 – 86.1 |
| Winter Humidity (%) | 78.1 | 10.6 | 47.4 – 86.3 |
| Summer NDVI* | 0.63 | 0.16 | 0.31 – 0.82 |
| Winter NDVI* | 0.29 | 0.16 | 0.02 – 0.75 |

\*NDVI – Normalized difference vegetation index

The host and region GLM models tested the same covariates across all gene segments and subtypes, individually. Overall, the internal gene segments held higher support for the inclusion of both host and region covariates as compared to the HA and NA subtype models (Figure 4). On average 20 of the 32 tested variables were supported for inclusion among the internal gene segments compared to five and seven supported variables among HA and NA subtypes, respectively. In the PDA sample, the H5 and N7 subtype models each supported only one variable across both host and region GLMs. Nonbreeding distribution overlap and migratory distance of the sink host species, as well as summer distribution overlap of the source region, winter temperature of both source and sink regions, winter precipitation of both source and sink regions, and winter humidity of source regions tended to have lower support among internal gene segment models. Overlap of host breeding distribution was supported across all gene segments and subtypes except H5 in both PDA and stratified samples and N6 in the PDA sample. Regional distance was also frequently supported across the gene segments and subtypes, with support in all but H10, N1, and N7 in both samples as well as H11 in the stratified sample. Summer temperature of the source region was supported in both samples for all internal gene segments as well as subtypes H1, H3, H4, H11, N2, and N3. Host genetic relatedness was supported in all internal gene segments and all but two NA subtypes, N7 and N9, yet there was no support for this variable among HA subtypes.

**Figure 4.**
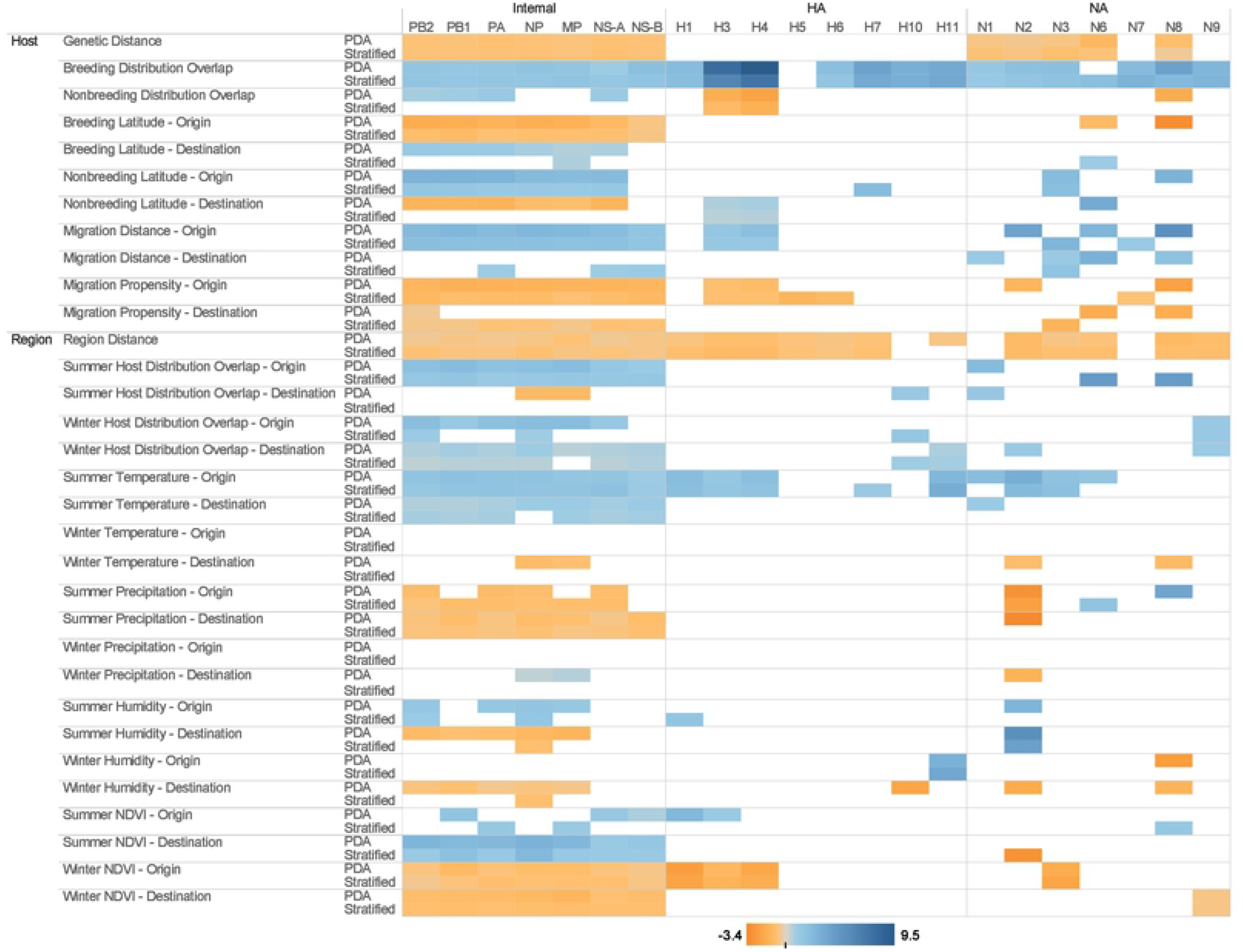
Heat map of conditional coefficient values for host and region generalized linear models of North American avian influenza discrete trait diffusion models. Conditional coefficient effect sizes are presented for each supported ecological variable across all gene segment and subtype datasets and both subsampling strategies (phylogenetic diversity analyzer (PDA) vs. stratified random sample). Only supported coefficients are displayed. Color darkness is proportional to the magnitude of the effect. Orange represents a negative correlation and blue represents a positive correlation.

The magnitude and direction of variable effect size differed among the various gene segments although most variables demonstrated the same directional effect across multiple gene segments. Among the variables supported in the models, 22 had the same directional effect (positive or negative) without regard to gene segment or subtype. Host variables which were consistently positively-associated with interspecies transmission included breeding distribution overlap, nonbreeding latitude and migration distance of the source host. Consistent negatively-associated host characteristics included genetic distance between host species and migration propensity of both the source and sink host species. Among North American regional geographic models, the proportion of avian hosts in source regions with summer distribution overlap, the proportion of avian hosts in sink regions with winter distribution overlap, both source and sink summer temperature measures, and source summer normalized difference vegetation index (NDVI) were positively associated with viral dispersal across all segments and subtypes in which the variables had support for inclusion. The following region characteristics were consistently negatively-associated with viral dispersal: distance between regions, summer precipitation of the sink region, and both winter NDVI measures. Several directional effects conflicted among subtypes and gene segments, including nonbreeding distribution overlap, nonbreeding latitude of the sink host, summer habitat overlap in the sink region, summer precipitation of the source region, winter precipitation of the sink region, summer humidity of the sink region, winter humidity of the source region, and summer NDVI of the sink region. These conflicts tended to be observed in variables with infrequent support, especially within the stratified sample.

## DISCUSSION

The presented analysis provides insight into the potential impact of ecological variables that influence AIV dispersal and diversity within North American wild birds. While the evolution and dispersal of AIV within North America has been previously examined (7,8,14,22,23), this study employs a discrete trait diffusion GLM to incorporate ecological data into such estimates. Using new and historical AIV sequences, we demonstrate that host and geographic characteristics are associated with viral movement among avian species and North American regions. Because AIV gene segments can be treated as independent hereditary particles, genetic similarities can be used to infer information regarding the ecological pressures experienced by viral populations. By estimating ecological models separately for each AIV gene segment, dispersal patterns and their associations with ecological characteristics can be tested independently and compared. Consistent support for a variable across multiple gene segments and subtypes, such as breeding distribution overlap and geographic distance between regions, suggest that these host habitat characteristics play an important role in the evolution and ecology of AIV. Although AIV hosts often migrate and potentially carry virus over long distances, a geographic distance effect can be noted: as the distance between two regions increases, the frequency of AIV transition decreases. The importance of proximity is reinforced by the consistent support of distribution overlap, particularly in the summer breeding season. A similar finding has been observed in bats, in which viral transmission of rabies virus was associated with host distribution overlap in North America (19). Species that have greater overlap during the breeding months tend to have a higher frequency of AIV transition due to a larger population of immunologically naïve juvenile hosts.

Another frequently supported host characteristic is the genetic distance or relatedness between two species, a characteristic that has been suggested to influence the rate of interspecies transmission of pathogens in general (24). Genetic relatedness may be a proxy for a suite of shared characteristics that would increase the likelihood of two hosts sharing a pathogen. For instance, viruses that infect multiple species are most likely targeting conserved molecular mechanisms, and related hosts will most likely have similar physiological responses (25). Furthermore, related species typically share similar ecology, i.e. breeding and feeding behavior or habitat, which can increase the likelihood of contact between the two species, a prerequisite for pathogen transmission. Experimental studies (26) and mathematical models (27) have shown that host relatedness is associated with successful host transition. Genetic distance, however, was not supported among any of the HA models. The HA models in general tended to have lower frequency of support for the included GLM models. This may suggest that the host immune pressure exerted on the HA supersedes influence of ecological determinants. In other words, because HA subtypes exist as a constellation of fitness peaks, these genes may be unable to provide information on ecological factors that affect viral transmission. Rather they are coerced by immune pressure to constantly accumulate mutations that provide fitness advantages to evade host immune systems.

Somewhat surprisingly, summer temperature of the originating region was positively associated with viral dispersal among regions in multiple gene segments. Environmental durability experiments (28,29) and AIV prevalence studies (11,12) have demonstrated evidence that colder temperatures increase risk of AIV infection due to environmental persistence of the virus. In contrast, our geographic model suggests that regions that are warmer on average during the summer are more likely to act as sources of the virus to other regions. It should be noted that causality cannot be established for this association. Proper interpretation of this result is that warmer regions are merely associated with viral dispersal, not that virus is more likely to arise from regions during summer. Summer temperature may be a proxy for other environmental or temporal characteristics. The effect of temperature on AIV dispersal can also be observed in the host models in which latitude of the breeding distribution was negatively associated with viral transitions between host species. In other words, species that breed farther south were more likely to act as sources of AIV diversity to other host species. Similarly, species that overwinter farther north were also more likely to act as sources of the virus. In corroboration with our model, one prevalence study (9) revealed an earlier thaw date of a location to be associated with higher AIV prevalence. Our results may be best explained by timing of breeding and migration rather than environmental persistence alone. Those locations that thaw first (i.e., are warmer in general) become available as breeding habitat sooner than regions farther north. Because breeding marks the influx of new, immunologically naïve juveniles, populations that breed earlier tend to become infected earlier, which may increase their capacity to serve as a source of virus to other hosts and locations.

As with analyses reliant on publicly available data, these results are limited by potential sampling bias of available surveillance and sequence data. As demonstrated, mallards markedly dominate the diffusion models as sources of virus to other species. Mallards are the most populous of the dabbling ducks and therefore are more frequently included in AIV surveillance, but they are often also the species with the highest prevalence of AIV (30). While one explanation for the estimation of mallards as frequent viral sources is their predominance in surveillance, the analysis methods were intended to limit the effects of sampling biases. Sequences collected prior to 2005 when sequencing efforts were irregular were not permitted to influence the discrete trait diffusion models. Further, by subsampling the datasets based on phylogenetics, we preserved the genetic diversity of the sequence data. The fact that mallards predominate in the PDA sample suggests that, as a primary reservoir species, mallards harbor a large diversity of AIV. A second subsampling technique (stratified random sample) was also performed in attempt to limit oversampling bias and increase the frequency of underrepresented hosts and regions. By comparing results between the two datasets, the influence of sampling schemes on the observed results can be approximated. Estimating the models across multiple gene segments and subtypes also allowed the host and regional proportions to vary, which is more apparent among the HA and NA subtypes. It should be noted that the magnitude of the effective population size across all segments tracked closely with the sample size of sequences included within the analysis. As the sample size was proportional to the number of available sequences that met inclusion criteria, the sample size may indicate the overall genetic diversity available for analysis, which would then be reflected in the estimated effective population size. This is supported by the use of phylogenetic diversity as a means to sub-sample the data, which ensures that both the full available sequences and the sampled sequences would have equivalent genetic diversity, unlike a simple random sample which, by chance, may remove some genetic diversity.

Although causality between ecologic factors and AIV diffusion cannot be inferred from this analysis, our results provide further evidence of the association of geographic and host characteristics with AIV diversity and dispersal. Continued AIV surveillance, especially in undersampled regions and hosts, provides valuable information on AIV evolution and diffusion. Furthermore, the inclusion of detailed environmental and host measures within AIV sequence databases will help add granularity to future models.

## MATERIALS AND METHODS

### Sequence data sets

Systematic avian influenza surveillance of wild birds has been performed in Alberta, Canada and Delaware Bay, New Jersey, United States since 1976 and 1985 respectively. Surveillance efforts, viral isolation, and genomic sequencing methods were performed as previously described (7). Newly sequenced genomes from 303 viral isolates were deposited in GenBank (Supplementary Table S11) and were aligned with publicly available AIV sequences from within North America between 1970 and 2016, which were downloaded from the Influenza Research Database (IRD; www.fludb.org) on March 26, 2018. Sequences with vague host (e.g., “avian,” “bird,” “duck,” etc.) or location (i.e., only country level data for the United States, Canada, or Mexico), more than 10% missing nucleotide sites, or isolated from domestic poultry were removed from the dataset. Internal gene segments PB2, PB1, PA, NP, and MP were aligned separately. Alignments of gene segments NS, HA, and NA were further subdivided: NS by allele group (A and B) and HA and NA by subtype. HA and NA subtypes with sparse representation (<250 sequences between 2005 and 2016) were excluded from the analysis (subtypes H2, H8, H9, H12, H13, H14, H16, N4, N5). Initial maximum likelihood phylogenetic trees were estimated using RAxML v8 (31) with a general time reversible nucleotide substitution model and gamma distribution of sites. TempEst v1.5 (32) was used to identify sequences with a rate of evolutionary divergence out of the expected bounds as compared to the remaining sequences in the dataset. This helps to identify poor quality sequences or viruses under unexpected evolutionary pressure. Eurasian lineages with little continued North American circulation or associated with highly pathogenic avian influenza viruses were removed from the dataset. For each sequence, host of origin was categorized based on host species. Location of origin was categorized based on United States National Oceanic and Atmospheric Administration historical climate region for United States isolates, province and territory for Canadian isolates, state for Mexican isolates, and country for Guatemalan isolates. Categories to be included in the discrete trait models were determined separately for the internal genes, HA subtypes, and NA subtypes. For internal genes, the 20 most common host species and regions (based on average rank among the segments) were chosen to be included in the discrete trait diffusion models. For both HA and NA subtypes, categories with greater than one sequence in a majority of the HA or NA subtypes were included in the models.

All sequences collected before 2005 were combined into a single category that was masked in the diffusion models to prevent the influence of inconsistent sampling and to focus diffusion summaries on the most recent years. To mitigate oversampling, two subsampling schemes were used: simple stratified random sampling and phylogenetic diversity-based sampling. In the simple random sample, sequences were stratified by region, host species, and year, and a maximum sample size of three sequences for each stratum were maintained in the dataset. Developed to help make economic decisions for conservation purposes, Phylogenetic Diversity Analyzer (PDA; http://www.cibiv.at/software/pda/) was used to select a subsample for each segment or subtype that maximized the represented genetic diversity (33). This process was weighted to prevent over-representation of samples before 2005 which, though diverse, were masked in the diffusion model. As PDA allows the user to select the desired sample size, the number of selected sequences was specified to match the stratified sample and ensure datasets were proportional.

### Phylogenetic analysis

Using ModelFinder algorithm (34) implemented in the program IQTree (http://www.iqtree.org/), the best fit nucleotide substitution model was determined. The empirical sets of phylogenetic trees were estimated under the same model assumptions for all sequence datasets in BEAST v1.10.4 (35). A general time reversible (GTR) nucleotide substitution model (36–38) with a 4-category gamma distribution of variation among sites and a proportion of invariant sites (39,40) was implemented with a lognormal uncorrelated relaxed molecular clock (41) (mean clock rate prior distribution: uniform 0 – 1, initial value =0.0033) and a constant coalescent population model (42,43) (population size prior distribution: lognormal distribution with mean = 50 and standard deviation = 50). At least four independent Markov chain Monte Carlo (MCMC) runs of 100 million state length and sampling every 10,000 states were performed. To ensure proper convergence and parameter mixing with an effective sample size (ESS) of at least 200, a minimum of 10% burn-in was removed. Non-convergent runs were discarded, larger burn-in percentages were removed, and additional MCMC runs were performed to achieve ESS > 200. Empirical tree sets were obtained by combining and resampling tree log files from non-discarded runs with LogCombiner to achieve a tree file length of at least 1,500 trees.

### Discrete trait diffusion models

With the ability to incorporate ecological and epidemiological metadata, the discrete trait diffusion model uses a continuous-time Markov chain as its basis to estimate the ancestral history of trait changes across a phylogenetic tree, in essence treating the trait as a characteristic that evolves over time (44,45). To investigate recent movement of AIV among avian hosts and North American regions, discrete trait diffusion models based on the empirical tree sets described above were estimated using BEAST v1.10.4. Estimating the posterior distribution of phylogenetic trees based on sequence data alone can be performed separately from the discrete trait diffusion models because the discrete traits represent only two sites (as opposed to the hundreds of nucleotide sites of a genetic sequence) at which the tree likelihood can be calculated. For this reason, the discrete trait model has an insignificant impact on phylogenetic estimation (16). Furthermore, this approach enables the inference of a single diffusion model across multiple empirical tree sets, allowing the genetic information from multiple gene segments to inform the model. Due to the high level of reassortment of gene segments within low pathogenic AIV in wild birds (5), each gene segment can be treated as an independent hereditary particle, providing separate evolutionary and ecological information within its phylogenetic history (7). Asymmetrical discrete trait diffusion models were estimated across empirical tree sets for the following: 1) each gene segment or subtype dataset individually, 2) all internal gene segments together, 3) all represented HA subtypes together, and 4) all represented NA subtypes together. Discrete host and geographic traits were specified as described above. Pre-2005 sequences and rare categories were masked from the discrete trait diffusion model, providing an estimate of viral transitions between common host species and regions between 2005 and 2016.

The discrete trait diffusion models were extended using a generalized linear model (GLM) to evaluate predictors associated with the discrete trait transition rates among host species and geographic regions. Using the transition rates as the outcome to a log-linear combination of covariate predictors, BEAST v1.10 estimates the GLM at each state in the MCMC simulation, integrating across the empirical phylogenetic tree space. Host diffusion predictors included genetic distance between species, habitat distribution overlap, migration distance, migration propensity, and latitudinal distribution. Genetic distance between species was calculated as the average patristic distance, represented as total evolutionary time in million years, across a sample of 1,000 phylogenetic trees estimated under a fossil-calibrated relaxed molecular clock (46). Habitat overlap, migration distance, migration propensity, and latitudinal distribution were summarized from BirdLife species range maps (47) using ArcGIS Pro software. Habitat distribution overlap was calculated as the percentage of a source host’s geographic distribution shared with that of a sink host. Migration distance was estimated by the difference between the mean breeding distribution latitude and the mean wintering distribution latitude (13). Migration propensity was estimated as the percentage of total summer distribution range considered to be migratory as opposed to resident (13). Latitudinal distribution was the average latitude for breeding and wintering ranges and served as an estimate of habitat temperature. Geographic diffusion predictors included distance between regions as well as summer and winter summaries of each of the following: average temperature, average precipitation, average relative humidity, average normalized difference vegetation index (NDVI), and proportion of included host species that reside in the region. All geographic variables were summarized between 2005 and 2016 and aggregated in ArcGIS Pro. Climatological data originated from the National Centers for Environmental Prediction North American Regional Reanalysis (48) provided by the National Oceanic and Atmospheric Administration Oceanic and Atmospheric Research Earth System Research Laboratory’s Physical Sciences Division, Boulder, Colorado, USA, from their website at https://www.esrl.noaa.gov/psd/. NDVI data originated from the Terra Moderate Resolution Imaging Spectroradiometer (MODIS) Vegetation Indices (MOD13A3) Version 6 (49). All covariates were log-transformed and standardized before inclusion in the GLM. Each discrete trait diffusion model and GLM were performed with at least three independent MCMC runs of 1 million chain length sampling every 100 states.

### Statistical analysis

For both the discrete trait diffusion model and the GLM, Bayesian stochastic search variable selection (BSSVS) was used to estimate the statistical support for diffusion rates and coefficients, respectively, by enabling the calculation of a Bayes factor (BF) (50). A larger BF indicates stronger support that the parameter (i.e., transition rate between two hosts or GLM coefficient) is non-zero. Due to the number of models tested, a stringent statistical cutoff was implemented, only allowing those with BF > 100 to signify statistical support. Median conditional transition rates, median conditional coefficients, 95% highest posterior density (HPD) intervals, and BF were calculated from BEAST posterior samples with personalized Python scripts using the PyMC3 package for Bayesian statistical modeling (51).

## DATA AVAILABILITY

Newly sequenced AIV nucleotide sequences have been deposited in GenBank with accession numbers provided in Supplementary Table S11.

## ACKNOWLEDGEMENTS

We would like to thank Karlie Woodard and Angela Danner for performing the RNA extraction as well as Nichola Hill for her insight into measures of avian phylogenetic diversity.

## SUPPLEMENTARY MATERIAL CAPTIONS

Figure S1. Evolutionary parameter estimation for North American avian influenza viruses of wild birds. Estimated parameters include A) time to most recent common ancestor (TMRCA), B) molecular clock rate, and C) effective population size. Parameters are compared across internal gene segments (blue), hemagglutinin gene subtypes (orange), and neuraminidase gene subtypes (purple) as well as between subsampling strategies, phylogenetic diversity-based sample (left, dark grey) and stratified random sample (right, light grey). Median values (black midline) indicated as well as the 95% highest posterior density (whiskers).

Figure S2. Host species temporal distribution of sampled North American avian influenza virus PB2 gene segment sequences, 2005 – 2016. Proportions of represented host species are compared between the phylogenetic diversity-based sample (PDA) and the stratified random sample (stratified).

Figure S3. Geographic region temporal distribution of sampled North American avian influenza virus PB2 gene segment sequences, 2005 – 2016. Proportions of represented regions are compared between the phylogenetic diversity-based sample (PDA) and the stratified random sample (stratified).

Figure S4. Heat map of supported viral transition rates among host species across avian influenza virus gene segments and subtypes. Colored cells represent the magnitude of the transition rate from the species in the first column (source) to the species in the second column (sink). White cells were transition rates that were not supported (Bayes factor < 100). Results from both subsampling strategies (phylogenetic diversity-based sample (PDA) and stratified random sample (stratified)) are presented for comparison.

Figure S5. Discrete trait diffusion models of North American avian influenza using a stratified random sample of genetic sequences. Host models (left) are presented for combined internal gene segment (A), hemagglutinin gene subtype (B), and neuraminidase gene subtype (C) models. Source host species on the left of the chord diagrams contribute viral diversity to sink host species on the right. The magnitude of the viral transition rate is proportional to the width of the band, and statistically supported rates darkened. Bands are colored by the host order of the source species (Charadriiformes – red; Anseriformes – blue). Similarly, geographic models (right) are summarized for combined internal gene segment (D), hemagglutinin gene subtype (E), and neuraminidase gene subtype (F) models. Arrow width is proportional to the magnitude of the transition rate, and only statistically supported rates are displayed. (AK – Alaska, AB – Alberta, BC – British Columbia, GT – Guatemala, MW – Midwest, NB – New Brunswick, NE – Northeast, NL – Newfoundland and Labrador, NRP – Northern Rockies and Plains, NS – Nova Scotia, NW – Northwest, OV – Ohio Valley, PE – Prince Edward Island, QC – Quebec, S – South, SE – Southeast, SON – Sonora, SW – Southwest, W – West)

Figure S6. Heat map of supported viral transition rates among geographic regions across avian influenza virus gene segments and subtypes. Colored cells represent the magnitude of the transition rate from the region in the first column (source) to the region in the second column (sink). White cells were transition rates that were not supported (Bayes factor < 100). Results from both subsampling strategies (phylogenetic diversity-based sample (PDA) and stratified random sample (stratified)) are presented for comparison.

Table S1. Demographic characteristics of 303 wild bird surveillance samples with newly sequenced avian influenza isolates, 2003 – 2016.

Table S2. Evolutionary parameters of avian influenza virus gene segments collected from North American wild birds between 1970 and 2016. Datasets were sampled so as to maintain the total phylogenetic diversity of the original publicly available sequence sample.

Table S3. Host and regional distribution of phylogenetic diversity-based subsample of influenza virus gene segments isolated from North American wild birds.

Table S4. Host and regional distribution of stratified subsample of influenza virus gene segments isolated from North American wild birds.

Tables S5 – S7. Host species transition rate matrix from combined internal gene model (Table S5), combined hemagglutinin subtype model (Table S6), and combined neuraminidase subtype model (Table S7). Median rates and 95% highest posterior density intervals are displayed for both subsampling strategies. Rates colored in blue are statistically supported (Bayes factor > 100). (ABD – American black duck, BUF – bufflehead, BWT – blue-winged teal, CAN – Canada goose, CIN – cinnamon teal, EMP – emperor goose, GAD – gadwall, GWF – greater white-fronted goose, GWG – glaucous-winged gull, GWT – green-winged teal, LAU – laughing gull, MAL – mallard, PIN – northern pintail, RED – redhead, RKN – red knot, RND – ring-necked duck, RUD – ruddy turnstone, SHO – northern shoveler, SND – sanderling, SNO – snow goose, WIG – American wigeon).

Tables S8 – 10. Geographic region transition rate matrix from combined internal gene model (Table S8), combined hemagglutinin subtype model (Table S9), and combined neuraminidase subtype model (Table S10). Median rates and 95% highest posterior density intervals are displayed for both subsampling strategies. Rates colored in blue are statistically supported (Bayes factor > 100). (AK – Alaska, ALB – Alberta, BCO – British Columbia, GUA – Guatemala, MW – Midwest, NBR – New Brunswick, NE – Northeast, NFL – Newfoundland and Labrador, RP – Northern Rockies and Plains, NSC – Nova Scotia, NW – Northwest, OV – Ohio Valley, PEI – Prince Edward Island, QUE – Quebec, S – South, SE – Southeast, SON – Sonora, SW – Southwest, W – West)

Table S11. Names and GenBank accession numbers of 303 newly sequenced AIV nucleotide sequences.

